# In macrophages fatty acid oxidation spares glutamate for use in diverse metabolic pathways required for alternative activation

**DOI:** 10.1101/2022.04.13.487890

**Authors:** Nikki van Teijlingen Bakker, Lea Flachsman, Gustavo E. Carrizo, David E. Sanin, Simon Lawless, Angela Castoldi, Lauar Monteiro, Agnieszka M. Kabat, Mai Matsushita, Fabian Haessler, Annette Patterson, Ramon Klein Geltink, David O’Sullivan, Erika L. Pearce, Edward J. Pearce

## Abstract

Fatty acid oxidation (FAO) is upregulated in IL-4-stimulated (alternatively activated) macrophages (M(IL-4)). We examined the effect of loss of function of the enzyme Cpt1a, which facilitates the entry of long chain fatty acids (FA) into mitochondria for FAO, on alternative activation. Expression of M(IL-4) markers ARG1, CD301 and RELMα, was impaired in tamoxifen-treated ERT2Cre x *Cpt1a*^*fl/fl*^ macrophages and in macrophages expressing shRNA targeting *Cpt1a (Cpt1a-*shRNA). In contrast, *Vavi*Cre x *Cpt1a*^*fl/fl*^ and *Lysm*Cre x *Cpt1a*^*fl/fl*^ M(IL-4) responded normally to IL-4. Reduced alternative activation due to Cpt1a loss of function was linked to decreased cellular pools of α-ketoglutarate, glutamate, and glutathione, diminished commitment of glucose carbon to serine/glycine synthesis, and decreased expression of genes in the Nrf2-oxidative stress response pathway. Consistent with this, reactive oxygen species were increased. Restoration of glutathione pools with N-acetyl cysteine normalized oxidative stress and allowed alternative activation in the face of *Cpt1a*-deficiency, pointing to a role for FAO in the control of ROS and as being important for alternative activation. In *Vavi*Cre x *Cpt1a*^*fl/fl*^ M(IL-4), glutamine uptake was increased, compensating for the loss of FAO to meet necessary metabolic demands, to allow alternative activation. The data indicate that macrophages are able to regulate glutamine metabolism to compensate for chronic disruption of FAO to meet metabolic needs.

## Introduction

Macrophages are tissue-dwelling innate immune cells that play diverse roles in homeostasis and inflammation. They are able to receive signals from immune system-intrinsic and extrinsic factors, in response to which they can assume functionally distinct activation states^1^. One defined macrophage activation state is that induced by STAT6-dependent signals delivered through the IL-4Rα by IL-4/IL-13. Macrophages “alternatively activated” (a term that stresses the contrast with the “classical activation” state induced by signals delivered through Toll like receptors^2^) in this way, express a set of genes including *Arg1* (arginase 1, ARG1), *Clec10a* (CD301) and *Retnla* (resistin-like molecule-alpha, RELMα), and are implicated in immunity against helminths^3^, wound-healing^4^, metabolic homeostasis^5^, the regulation of classical inflammation, and cancer progression and metastasis^1,6^.

Macrophage activation is accompanied by, and dependent upon, cell-intrinsic metabolic changes which reflect distinct metabolic requirements for different activation processes^7^. For example, macrophages alternatively activated in response to IL-4 (M(IL-4)) exhibit increased oxidative phosphorylation (OXPHOS)^8,9^, fueled by increased fatty acid oxidation (FAO)^10,11^. FAO is dependent on carnitine palmitoyltransferase 1a (Cpt1a), which produces long chain acyl-carnitines from long chain fatty acids (Lc-FAs), that can then be transported across the outer membrane of the mitochondria by the carnitine-acylcarntine transporter Slc25a20. Cpt2 mediates the removal of carnitine from acyl-carnitines, allowing long chain acyl-CoAs to cross the inner mitochondrial membrane and enter cycles of FAO^12^ in the matrix. FAO, and additionally the uptake of FA and their generation from triacylglycerol by lipolysis, are all increased in macrophages stimulated with IL-4 and interference with different steps in these processes has been reported to inhibit M(IL-4) function *in vitro* and/or *in vivo*^11,13–16^. To some extent, conclusions on the importance of FA catabolism in alternative activation have arisen from findings that etomoxir, a small molecule inhibitor of FAO, is able to inhibit M(IL-4) activation^10,11^. However, in recent reports IL-4-induced alternative activation was reported to proceed normally in *LysM*Cre x *Cpt1a*^*fl/fl*^ and *LysM*Cre x *Cpt2*^*fl/fl*^ bone marrow derived macrophages^17,18^, and effects of etomoxir were assigned to off-target effects on the intracellular CoA pool^18^.

In addition to FAO, recent work has revealed important roles in alternative activation for the entry of glucose and glutamine-derived carbon into the TCA cycle^7,14^. Pharmacologic inhibition of the catabolism of glucose and glutamine have both been reported to inhibit alternative activation^10,11,15,19,20^. Glutamine is additionally important because it is a precursor of the TCA cycle intermediate α-ketoglutarate (αKG), which can promote the Jmjd3-dependent demethylation of repressive H2K27me3 marks around M(IL-4) genes^20^. Glucose has been shown to be important for alternative activation because of its role as a carbon source for the production of citrate as a source of acetyl-CoA, which is a donor for histone acetylation to allow the expression of M(IL-4) genes^14^. In contrast to the situation for glucose and glutamine, the molecular basis for a role of FA in alternative activation is less well understood, while FAO itself may play additional roles in alternative activation that remain to be determined.

To explore the role of FAO in M(IL-4) in greater detail, we examined the role of Cpt1a using a panel of distinct genetic loss of function models. Acute *Cpt1a*-shRNA induced knock down, or tamoxifen induced deletion of *Cpt1a* in ERT2Cre x *Cpt1a* ^*fl/fl*^ macrophages *in vitro*, resulted in impaired alternative activation after IL-4 stimulation. In contrast, alternative activation was unaffected by deletion of *Cpt1a* in *Lysm*Cre x *Cpt1a* ^*fl/fl*^ or *Vavi*Cre x *Cpt1a* ^*fl/fl*^ macrophages. Consistent with a central role for Cpt1a in FAO in macrophages, acyl-carnitines and FAO were reduced across loss of function models. This led to diminished pools of TCA cycle intermediates, including αKG, in *Cpt1a*-shRNA and ERT2Cre x *Cpt1a* ^*fl/fl*^ M(IL-4), but not in *Vavi*Cre x *Cpt1a* ^*fl/fl*^ M(IL-4). Rather, these latter cells were able to maintain pools of TCA cycle intermediates through increased use of glutamine-derived carbon. In *Cpt1a*-shRNA and ERT2Cre x *Cpt1a* ^*fl/fl*^ M(IL-4), dysregulated glutamate metabolism linked to declines in serine/glycine synthesis resulted in diminished glutathione levels and increased levels of reactive oxygen species (ROS), which were not apparent in *Vavi*Cre x *Cpt1a* ^*fl/fl*^ M(IL-4). The addition of N-acetyl cysteine (NAC) to *Cpt1a*-shRNA and ERT2Cre x *Cpt1a* ^*fl/fl*^ M(IL-4) alleviated oxidative stress and restored the ability of IL-4 stimulated macrophages to express markers of alternative activation.

## Results

### The effect of the loss of Cpt1a function on alternative activation

We used a variety of approaches to explore the effect of Cpt1a loss of function on the ability of macrophages to become alternatively activated in response to IL-4. Using flow cytometry, we found that the expression of the alternative activation markers ARG1, CD301 and RELMα was significantly impaired in tamoxifen-treated ERT2Cre x *Cpt1a*^fl/fl^ bone marrow derived macrophages (BMDM) compared to in tamoxifen-treated *Cpt1a*^fl/fl^ WT BMDM (Fig. 1a, S1a; model-specific controls are referred to as “WT”, but are defined in detail in Fig. S1a, d, h, for each model utilized). Consistent with reduced ARG1 expression, cellular arginine levels were increased in IL-4 treated ERT2Cre x *Cpt1a*^fl/fl^ BMDM (Fig. S1b). Further, we examined TIM4^+^ peritoneal macrophages (pMac) isolated from ERT2Cre x *Cpt1a*^fl/fl^ mice (Fig. S1c), treated with tamoxifen *in vivo*, and found that while their frequency *in vivo* was unaffected by *Cpt1a* deletion (Fig. S1c), their ability to express ARG1, CD301 and RELMα in response to IL-4 *in vitro* was also impaired (Fig. 1b).

**Figure 1.**
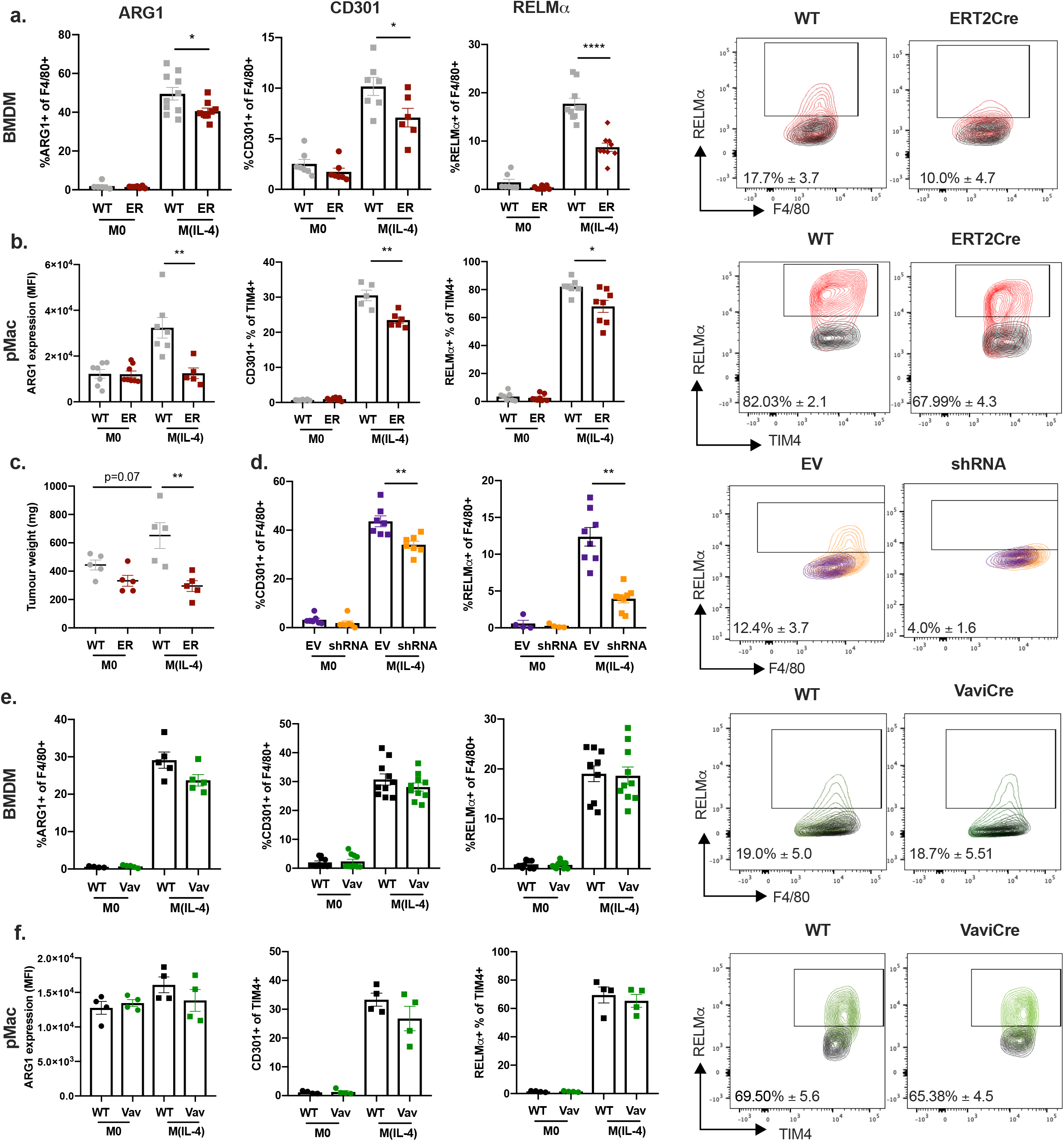
The effect of loss of Cpt1a function on the response of macrophages to IL-4. a. Expression of ARG1, CD301 and RELMα as percentage of F4/80+ cells in *Cpt1a*^fl/fl^ (WT, grey) or ERT2Cre x *Cpt1a*^fl/fl^ (ER, red) resting (M0) or IL-4 stimulated (M(IL-4)) BMDM, as measured by flow cytometry. Right hand panels show staining for RELMα, numbers signify mean RELMα % ± SEM of WT or ERT2Cre (ER) M(IL-4). b. Expression of indicated markers as median fluorescence intensity (MFI) or percentage positive of WT or ER M0 or M(IL-4) TIM4+ pMacs, as measured by flow cytometry. Right hand panels show staining for RELMα, numbers signify mean RELMα % ± SEM of WT or ER M(IL-4). c. Weight (mg) of day 14 B16 melanoma tumours initiated by co-injection of tumor cells plus WT or ER M0 or M(IL-4) BMDM. d. Expression of CD301 and RELMα as percentage of F4/80+ cells in EV-shRNA (EV, purple) or *Cpt1a*-shRNA (shRNA, orange) transduced M0 or M(IL-4), as measured by flow cytometry. Right hand panels show staining for RELMα, numbers signify mean RELMα % ± SEM of EV or shRNA M(IL-4). e. Expression of ARG1, CD301 and RELMα as percentage of F4/80+ cells in *Cpt1a*^fl/fl^ (WT, black) or *Vavi*Cre x *Cpt1a*^fl/fl^ (Vav, green) M0 or M(IL-4), as measured by flow cytometry. Right hand panels show staining for RELMα, numbers signify mean RELMα % ± SEM of WT or *Vavi*Cre M(IL-4). f. Expression of ARG1, CD301 and RELMα as median fluorescence intensity (MFI) of WT or Vav M0 or M(IL-4) TIM4+ pMacs, as measured by flow cytometry. Right hand panels show staining for RELMα, numbers signify mean RELMα % ± SEM of WT or *Vavi*Cre M(IL-4). Data represent mean ± SEM from 10 (a), 7 (b), 6 (c), 8 (d), 5 (e), 4 (f) biological replicates. Data represent three (a, d, e) or two (b, f) independent experiments. (*p < 0.05, **p < 0.01, ****p < 0.0001).

Alternatively activated macrophages mediate a range of processes *in vivo*, including the promotion of tumor growth^1,6^. We asked whether the latter was affected by Cpt1a deletion. For these studies, we employed a BMDM, B16 melanoma co-injection model, since it has been shown that co-injecting M(IL-4) with B16 melanoma cells leads to exacerbated tumour growth^21^. We found that, as expected, tumor growth was promoted when WT M(IL-4) *versus* M0 macrophages were co-injected with B16 melanoma cells^21–23^ (Fig. 1c). However, ERT2Cre x *Cpt1a*^fl/fl^ M(IL-4) failed to promote tumour growth to the extent observed for WT M(IL-4) (Fig. 1c).

Our findings with ERT2Cre x *Cpt1a*^fl/fl^ BMDM are consistent with previous reports that *Cpt1a*-siRNA suppresses alternative activation^11^. Indeed, when we targeted *Cpt1a* using *Cpt1a*-shRNA (Fig. S1d), we found that, as in ERT2Cre x *Cpt1a*^fl/fl^ BMDM, IL-4 induced alternative activation was negatively affected, with CD301 and RELMα expression significantly diminished (Fig. 1d), and cellular arginine levels increased (Fig. S1e), compared to in control empty vector (EV) shRNA BMDM stimulated with IL-4. However, none of these data are consistent with findings from *Lysm*Cre x *Cpt1a*^fl/fl^ mice, where alternative activation was reported to be unaffected by *Cpt1a* deletion^18^. We therefore directly examined IL-4-induced alternative activation in *Lysm*Cre x *Cpt1a*^fl/fl^ macrophages. Further, we assessed the effect of Cpt1a loss of function in *Vavi*Cre x *Cpt1a*^fl/fl^ macrophages (Fig. 1e, S1h). Cpt1a expression was significantly reduced in each of these models, although differences between systems were apparent with *Vavi*cre resulting in the most robust reduction (Fig. S1f, h). We found that in response to IL-4, the expression of CD301 and RELMα in *Lysm*Cre x *Cpt1a*^fl/fl^ BMDM was equivalent to that in WT cells (Fig. S1g), as previously published^18^. Similarly, expression of ARG1, CD301 and RELMα in IL-4-stimulated *Vavi*Cre x *Cpt1a*^fl/fl^ BMDM and pMacs was similar to that in WT IL-4 stimulated controls (Fig. 1e,f), as were cellular levels of arginine (Fig. S1i).

Taken together, these data show that the effect of *Cpt1a* deletion on the ability of macrophages to respond to IL-4 is dependent on the approach used to reduce Cpt1a expression. Interestingly, the effect on alternative activation did not correlate with the extent to which Cpt1a expression was decreased, since optimal Cpt1a deletion occurred in the *Vavi*Cre x *Cpt1a*^fl/fl^ model, and yet in response to IL-4, macrophages from these mice expressed alternative activation markers equivalently to WT macrophages. We postulate that the more acute loss of function in the ERT2Cre x *Cpt1a*^fl/fl^ and *Cpt1a*-shRNA model *versus* the chronic deletion in *Lysm*Cre x *Cpt1a*^fl/fl^ and *Vavi*Cre x *Cpt1a*^fl/fl^ models may be a differentiating feature.

### Reduction in Cpt1a expression leads to reduced FAO capacity

We sought to understand the metabolic impact of *Cpt1a* deletion. Considering the role of Cpt1a in Lc-FAO we began by examining the effect of *Cpt1a* deletion on FA-fueled OXPHOS. We took advantage of the fact that the addition of palmitate is expected to increase oxygen consumption rates (OCR) commensurate with the capacity of the cells to perform Lc-FAO. We found that palmitate increased baseline and maximal OCR in WT M(IL-4) macrophages, but not M0 macrophages, and that this increase was absent in both *Vavi*Cre x *Cpt1a*^fl/fl^ and ERT2Cre x *Cpt1a*^fl/fl^ M(IL-4) macrophages, consistent with the inability of both cell types to use palmitate to fuel respiration (Fig. 2a, b). These findings indicated that deletion of *Cpt1a* prevented M(IL-4) from utilizing palmitate to promote OXPHOS. To explore this further, we used mass spectrometry to measure IL-4 induced changes in palmitate metabolism. We found that, consistent with increased OXPHOS in M(IL-4) (Fig. 2a, b), the overall cellular pools of citrate, glutamate and malate were increased in IL-4-stimulated compared to resting macrophages (Fig. S2a). Consistent with the role of Cpt1a in FAO, ^13^C-palmitate incorporation into palmitoyl-carnitine pools, and overall cellular pools of parmitoyl carninite in IL-4 stimulated macrophages, were significantly diminished as a result of loss of Cpt1a function in *Vavi*Cre x *Cpt1a*^fl/fl^, ERT2Cre x *Cpt1a*^fl/fl^ and *Cpt1a*-shRNA BMDM (Fig. 2c, S2b, c, d).

**Figure 2.**
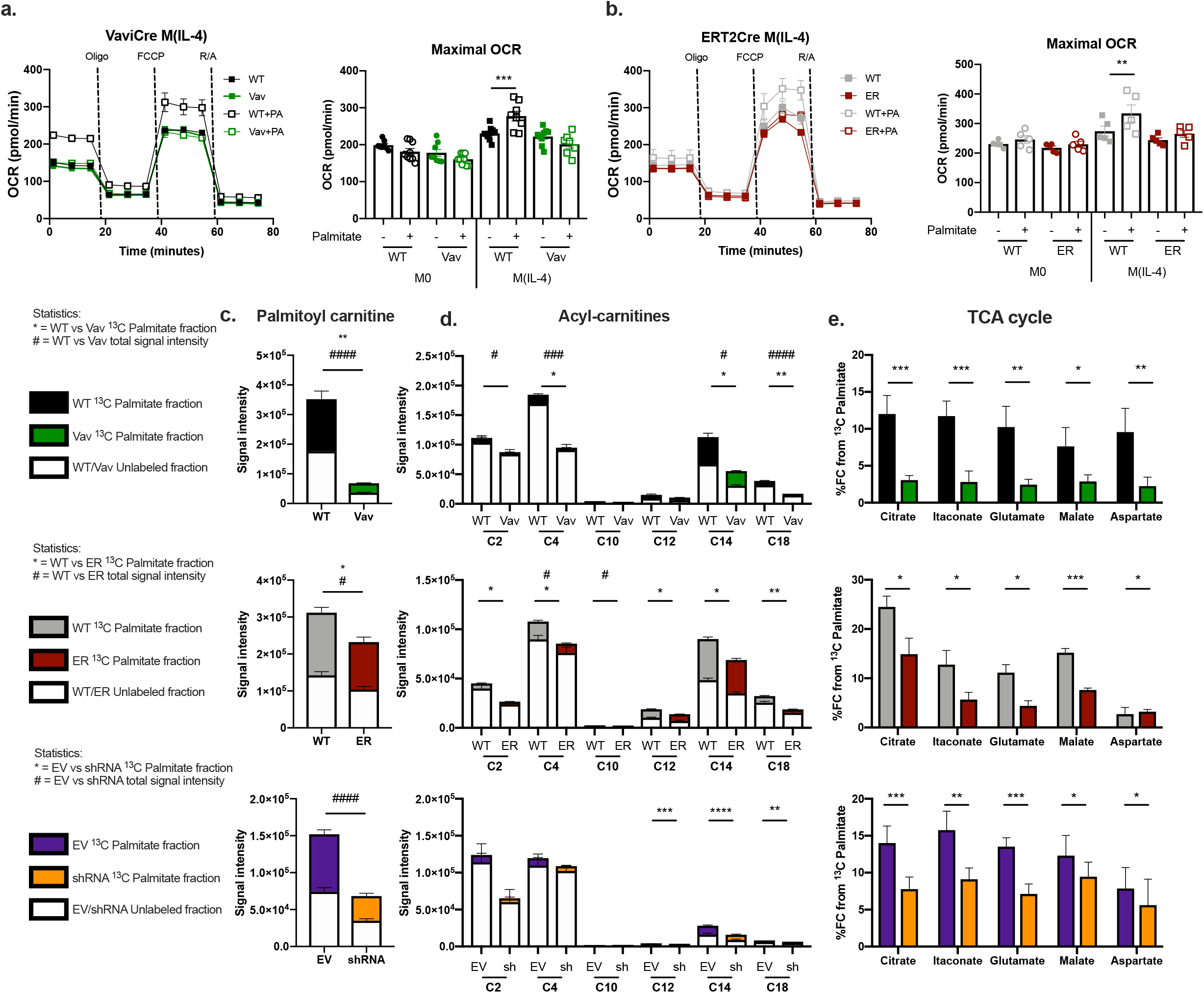
Reduction in Cpt1a expression leads to reduced FAO capacity. a. Extracellular flux analysis of oxygen consumption rate (OCR) of M0 or M(IL-4) BMDM from *Cpt1a*^fl/fl^ (WT, black) or *Vavi*Cre x *Cpt1a*^fl/fl^ (Vav, green) with or without 100 μM Palmitate (PA). After incubation with or without palmitate in XF media, cells were sequentially treated with oligomycin (oligo), FCCP, and rotenone plus antimycin (R/A). Maximal OCR is the average of the three measurements after FCCP injection. Error bars represent mean ± SEM, from 10 biological replicates. b. Extracellular flux analysis of OCR of M0 or M(IL-4) BMDM from *Cpt1a*^fl/fl^ (WT, grey) or ERT2Cre x *Cpt1a*^fl/fl^ (ER, red) with or without 100 μM Palmitate. After incubation with or without palmitate in XF media, cells were sequentially treated with oligo, FCCP, and R/A. Maximal OCR is the average of the three measurements after FCCP injection. Error bars represent mean ± SEM, 6 biological replicates. c. ^13^C-Palmitate LC-MS tracing into palmitoyl carnitine in WT or EV (purple), or Vav, ER and *Cpt1a*-shRNA (shRNA, orange) BMDM after stimulation with IL-4. d. ^13^C-Palmitate LC-MS trace into acyl-carnitines of different carbon chain sizes (C2-C18) in WT or EV and Vav, ER or shRNA BMDM after stimulation with IL-4. e. ^13^C-Palmitate LC-MS trace into citrate, itaconate, glutamate, malate and aspartate in WT or EV, or Vav, ER and shRNA BMDM after stimulation with IL-4. White section of the bar is the unlabeled fraction and coloured section of the bar is ^13^C-labeled fraction of the total metabolite pool for WT (black), Vav (green), WT (grey), ER (red), EV (purple) and shRNA (orange). Data represent mean ± SEM mean ± SEM, from 4 (Vav), 3 (ER) and 4 (shRNA) biological replicates. * represents statistics on ^13^C fractional contribution and # represents the statistics on the intracellular metabolite pools. A, b, c, d, e. Data represent two independent experiments. (*/#p < 0.05, **/##p < 0.01, ***/###p<0.001, ****/####p < 0.0001).

FA entry into the mitochondrial matrix from the intermembrane space is facilitated by Cpt2. We found that palmitoyl carnitine accumulated in *Cpt2*-shRNA M(IL-4) cultured with added palmitate, consistent with ongoing production of this acyl-carnitine in a situation where the enzyme required to remove carnitine, Cpt2, was absent (Fig. S2d, e). Consistent with the inability of *Cpt2*-shRNA M(IL-4) to oxidize Lc-FA, these cells, like *Cpt1a*-shRNA M(IL-4), exhibited impaired alternative activation (Fig. S2f).

Broad analysis of acyl carnitines in ^13^C-palmitate labeled M(IL-4) BMDM revealed palmitate carbon in long-, medium-, and short-chain acyl-carnitine pools, consistent with the breakdown of palmitate by FAO (Fig. 2d). Loss of Cpt1a function in *Vavi*Cre x *Cpt1a*^fl/fl^, ERT2Cre x *Cpt1a*^fl/fl^ and *Cpt1a*-shRNA BMDM, led to generally diminished ^13^C-palmitate contribution and reduced pools of short-, medium-and long-chain acyl carnitines (Fig. 2d). This was associated with reduced palmitate carbon incorporation into citrate, glutamate and malate across loss of function models (Fig. 2e).

### Glutamine is the preferred carbon source to maintain glutamate levels in M(IL-4)

Consistent with increased TCA cycle metabolites in M(IL-4) compared to M0, as well as the role of FAO in fueling the TCA cycle, reduced FAO due to Cpt1a loss of function was associated with a reduction in total citrate and glutamate pools in IL-4-stimulated ERT2Cre x *Cpt1a*^fl/fl^, *Cpt1a*-shRNA BMDM (Fig. 3a, b), and *Cpt2*-shRNA BMDM (Fig. S3a). However, this was not the case in *Vavi*Cre x *Cpt1a*^fl/fl^ M(IL-4) macrophages (Fig. 3a, b). Taken together, our data indicate that reductions in cellular pools of citrate and glutamate are metabolic characteristics of macrophages in which Cpt1a loss of function is associated with diminished potential for alternative activation.

**Figure 3.**
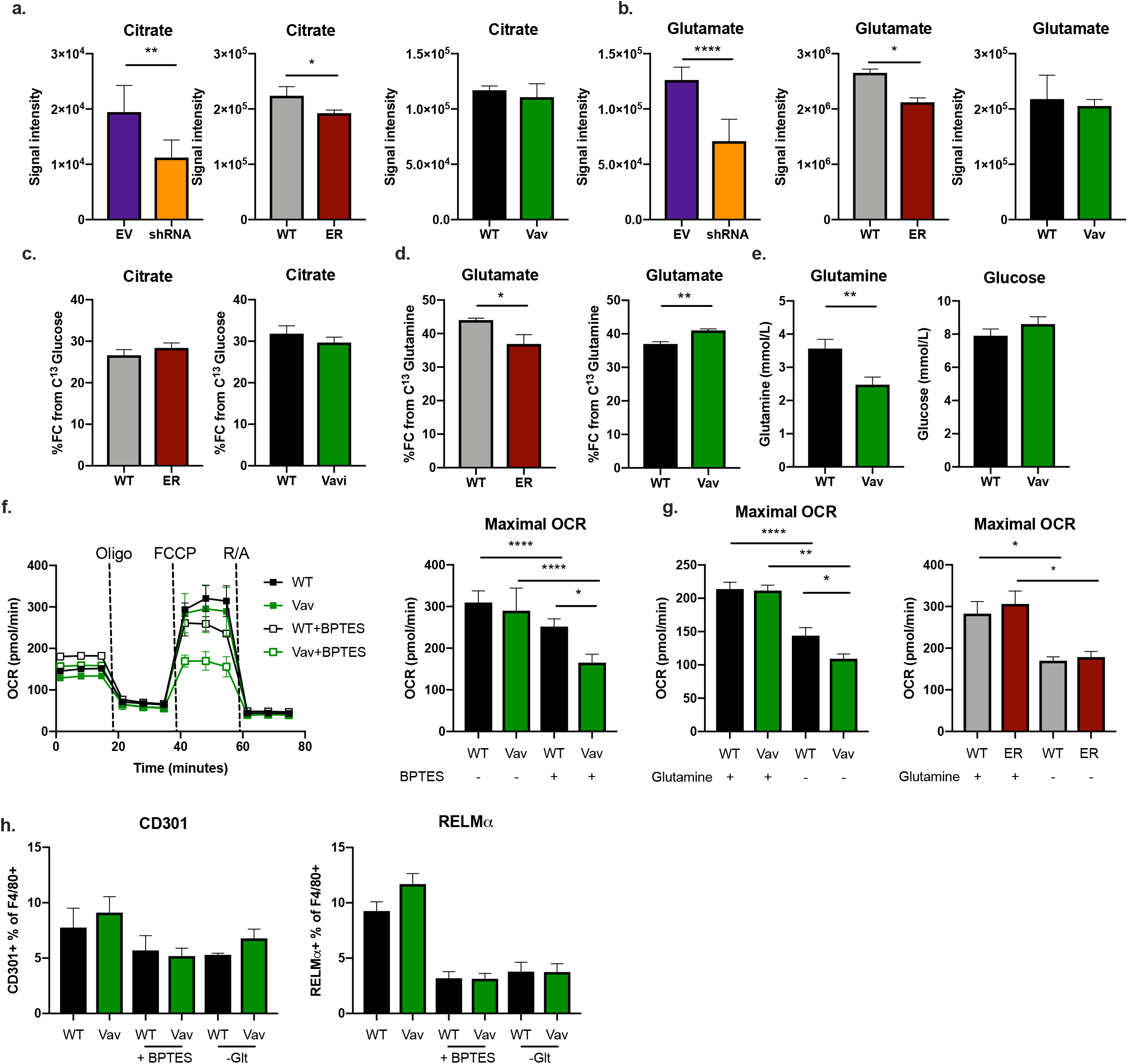
Glutamine is the preferred carbon source to maintain glutamate levels in M(IL-4) A.Signal intensity of citrate in empty vector (EV, purple), *Cpt1a*-shRNA (shRNA, orange), wildtype (WT, grey), ERT2Cre x *Cpt1a*^fl/fl^ (ER, red), wildtype (WT, black) and *Vavi*Cre x *Cpt1a*^fl/fl^ (Vav, green) as measured by LC-MS. b. Signal intensity of glutamate in EV vs shRNA, WT vs ER and WT vs Vav as measured by LC-MS. c. %^13^C-glucose fractional contribution (FC) to citrate in WT vs ER and WT vs Vav BMDM stimulated with IL-4, measured by LC-MS. d. %^13^C-glutamine fractional contribution (FC) to glutamate in WT vs ER and WT vs Vav BMDM stimulated with IL-4, measured by LC-MS. e. Glutamine and glucose concentrations in the supernatants of WT or Vav BMDM stimulated with IL-4, measured by bioanalyzer. f. Extracellular flux analysis of OCR of WT or Vav BMDM stimulated IL-4 with or without 10 μM BPTES. Maximal OCR is the average of the three measurements after FCCP injection. g. Extracellular flux analysis of OCR in the presence or absence of 1 mM glutamine of WT and Vav or WT and ER BMDM stimulated with IL-4. Maximal OCR is the average of the three measurements after FCCP injection. h. Expression of RELMα and CD301 as percentage of F4/80+ cells in WT or Vav BMDM, as measured by flowcytometry, in cells stimulated with IL-4 with or without BPTES or Glutamine (-Glt). Data represent mean ± SEM mean ± SEM from 3 (ER a, b, c, d), 4 (shRNA, Vav a, b, c, d), 8 (e), 3 (f, g, h) biological replicates representing two (a, b, c, d, e) or three (f) independent experiments. (*p < 0.05, **p < 0.01).

We hypothesized that in *Vavi*Cre x *Cpt1a*^fl/fl^ M(IL-4) the maintenance of key metabolic intermediates through alternative fuel use might compensate for reduced FAO to support alternative activation. To explore this possibility, we specifically analyzed the use of glucose and glutamine, which have been shown previously to be critical for alternative activation^10,11,15,19,20^. Glucose enters the TCA cycle through citrate, while glutamine can replenish the TCA cycle through αKG produced from glutamate during glutaminolysis. We found that the incorporation of ^13^C glucose into citrate was equivalent in *Vavi*Cre x *Cpt1a*^fl/fl^ and ERT2Cre x *Cpt1a*^fl/fl^ BMDM (Fig. 3c), but that incorporation of ^13^C glutamine into glutamate, was significantly greater in *Vavi*Cre x *Cpt1a*^fl/fl^ M(IL-4) but not ERT2Cre x *Cpt1a*^fl/fl^ M(IL-4), compared to in WT M(IL-4) (Fig. 3d). In line with these results, glutamine, but not glucose, was depleted to a greater extent in cultures of IL-4-stimulated *Vavi*Cre x *Cpt1a*^fl/fl^ BMDM (Fig. 3e). The importance of glutamine in fueling OXPHOS in the absence of FAO in these cells was revealed by extracellular flux analysis in the presence of the glutaminolysis inhibitor BPTES, which caused a marked decrease in spare respiratory capacity (SRC) and maximal OCR in IL-4 stimulated *Vavi*Cre x *Cpt1a*^fl/fl^ compared to WT BMDM (Fig. 3f). Glutamine depletion from tissue culture medium also caused a comparatively greater decrease in maximal OCR in *Vavi*Cre x *Cpt1a*^fl/fl^ M(IL-4) than in WT M(IL-4) (Fig. 3g), suggesting that glutamine was being used preferentially in the *Vavi*Cre x *Cpt1a*^fl/fl^ versus WT cells. The critical role for glutamine in alternative activation was illustrated by the finding that glutamine depletion and BPTES treatment markedly inhibited the expression of CD301 and RELMα in both WT and *Vavi*Cre x *Cpt1a*^fl/fl^ M(IL-4) (Fig. 3f).

Our data indicate that IL-4-stimulated *Vavi*Cre x *Cpt1a*^fl/fl^ BMDM are more capable than ERT2Cre x *Cpt1a*^fl/fl^ BMDM of using glutamine carbon to fuel the TCA cycle, and we speculate that in the *Vavi*Cre x *Cpt1a*^fl/fl^ cells this is sufficient to allow IL-4-induced alternative activation in the absence of FAO. We hypothesized that chronic, compared to more acute, deletion of *Cpt1a* allows cells to adapt in this way. We sought evidence to support this hypothesis from a distinct system. We compared the effects of chronically deleting *Mpc2*, a critical component of the mitochondrial pyruvate carrier, with acutely inhibiting this carrier with UK5099, which has previously been shown to negatively affect alternative activation^19^. We found that MPC2 was deleted in *Lysm*-cre x *Mpc2*^fl/fl^ BMDM (Fig. S3b), but contrary to the situation in UK5099-treated cells, the *Lysm*Cre x *Mpc2*^fl/fl^ BMDM responded normally to stimulation with IL-4 (Fig. S3c, d). Both IL-4 stimulated *Lysm*Cre x *Mpc2*^fl/fl^ and UK5099 treated BMDM incorporated less carbon from glucose into citrate and glutamate (Fig. S3e), however in *Lysm*Cre x *Mpc2*^fl/fl^ BMDM, but not UK5099-treated M(IL-4), the use of glutamine carbon increased and allowed the maintenance of TCA cycle metabolite pools (Fig. S3f, g, h). Taken together, our findings support the view that increased glutamine metabolism can be used by macrophages to compensate for the chronic loss of the ability to catabolize FA or glucose to support the TCA cycle and OXPHOS, and that this allows metabolic flexibility to support alternative activation.

### Acute loss of Cpt1a dysregulates glutamate metabolism and gene expression

Our data showed that increases in glutamine metabolism can compensate for reductions in the catabolism of FA or glucose to support alternative activation, and raised the possibility that this is related to the maintenance of adequate TCA cycle metabolite pools. αKG, a TCA cycle intermediate specifically implicated in alternative activation^20^, is produced from isocitrate by isocitrate dehydrogenase 2, but also directly from glutamine, through glutamate, during glutaminolysis^24^. We found that αKG pools were decreased in *Cpt1a-*shRNA and ERT2Cre x *Cpt1a*^fl/fl^ M(IL-4), but not in *Vavi*Cre x *Cpt1a*^fl/fl^ M(IL-4) (Fig. 4a, Fig. S4a). Functionally, αKG plays a role in epigenetically reprogramming alternative activation genes by promoting the demethylation of H3K27 by Jmjd3^20^. Consistent with these effects, chromatin accessibility around transcription start sites was diminished in *Cpt1a-*shRNA BMDM compared to EV-shRNA BMDM in M0 and to a significantly greater degree in M(IL-4) (Fig. 4b).

**Figure 4.**
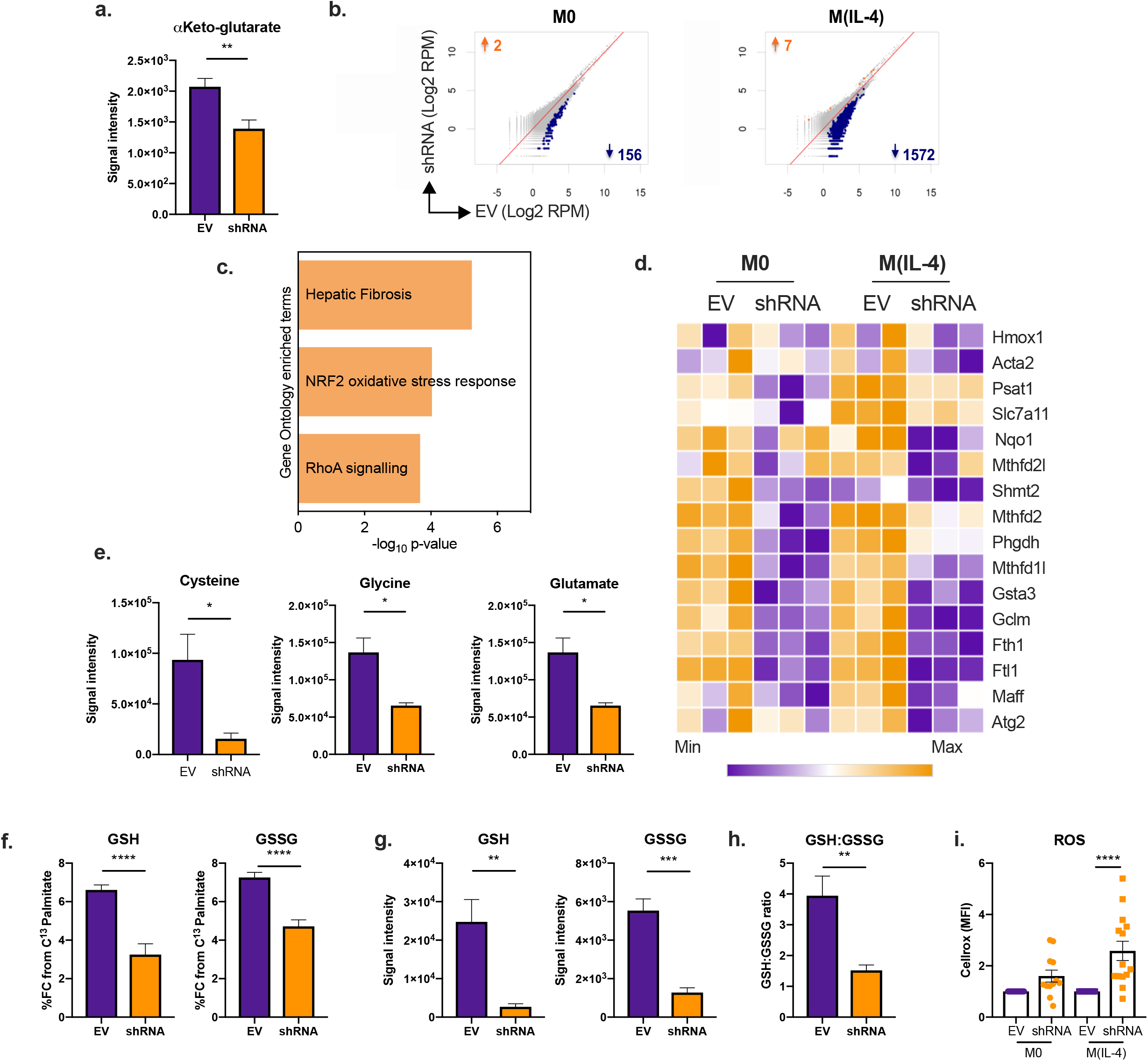
Acute loss of Cpt1a dysregulates glutamate metabolism and increases oxidative stress. a. Intracellular α-KG levels, as measured by LC-MS in empty vector (EV, purple) and *Cpt1a*-shRNA (shRNA, orange) BMDM stimulated with IL-4. b. Regions of accessible chromatin around the transcription start site that were significantly closed (↓) or opened (↑), as determined by ATAC-seq, in *Cpt1a*-shRNA compared to empty vector in resting (M0) or IL-4 stimulated (M(IL-4)) BMDM. c. Top 3 regulated pathways in shRNA compared to EV M(IL-4) as determined by Ingenuity Pathway Analysis (IPA) sorted by p-value and determined by the gene ontology enriched terms. d. Heatmap of genes indicated as NRF2 oxidative stress response genes that were up/down-regulated significantly (p<0.01) more than 2-fold in shRNA M(IL-4) compared to EV M(IL-4). e. Intracellular metabolite levels of cysteine, glycine and glutamate in EV and shRNA M(IL-4), as measured by LC-MS. f. ^13^C-Palmitate fractional contribution (FC) to reduced glutathione (GSH) and oxidized glutathione (GSSG) in EV and shRNA M(IL-4) as determined by LC-MS. g. Signal intensity of GSH and GSSG metabolite pools as determined by LC-MS in EV and shRNA M(IL-4). h. Ratio of GSH to GSSG of global metabolite pools in EV and shRNA M(IL-4). i. Cellular reactive oxygen species (ROS) determined by cellrox median fluoresence intensity (MFI) of shRNA normalised to EV M0 or M(IL-4). Data represent mean ± SEM mean ± SEM from 4 (a-h) and 12 (i) biological replicates. Data represent two (a, e-i) independent experiments. (**p < 0.01, ***p<0.001, ****p < 0.0001).

Alterations in chromatin accessibility would be expected to result in transcriptional changes. Ingenuity pathway analysis of RNAseq data from control and IL-4 stimulated *Cpt1a*-shRNA and EV-shRNA BMDM revealed that one of the most regulated pathways associated with loss of Cpt1a function was the NRF2 oxidative stress response (Fig. 4c). Genes in this pathway were significantly downregulated more than two-fold as a result of Cpt1a loss of function in IL-4 stimulated cells (Fig. 4d) and the results were recapitulated in *Cpt2*-shRNA BMDM (Fig. S4b). Amongst these genes were *Phgdh, Psat1, Shmt2, Mthfd2, Mthfd2l, Slc7a11, Gsta3 and Gclm*, which encode key enzymes in glutathione synthesis. Some of these genes, including *Phgdh* and *Slc7a11*, were amongst those in M(IL-4) in which chromatin became less accessible as a result of Cpt1a loss of function (Fig. S4c). In parallel, we found that the overall cellular pools of cysteine, glycine and glutamate, the essential building blocks of glutathione, were significantly reduced in *Cpt1a*-shRNA M(IL-4) (Fig. 4e).

Glutamate is integral to the synthesis of glutathione, which is critical for reducing ROS^25^. The use of palmitate for glutamate synthesis has been reported previously^26^, and consistent with this, we found that ^13^C-palmitate could be traced into reduced and oxidized glutathione (GSH and GSSG respectively), but that the labeling of both forms in IL-4 stimulated cells was diminished in *Cpt1a-*shRNA compared to EV-shRNA BMDM (Fig. 4f), as well as in ERT2Cre x *Cpt1a*^fl/fl^ compared to WT BMDM (Fig. S4d), and IL-4 stimulated *Vavi*Cre x *Cpt1a*^fl/fl^ BMDM compared to WT BMDM (Fig. S4e). Nevertheless, our data indicated that overall pools of both GSH and GSSG were greatly diminished in *Cpt1a-*shRNA M(IL-4) compared to EV-shRNA M(IL-4) (Fig. 4g), and the ratio of reduced to oxidized glutathione (GSH:GSSG) was decreased in IL-4 stimulated *Cpt1a-*shRNA (Fig. 4h), as well as in ERT2Cre, but not in *Vavi*Cre x *Cpt1a*^fl/fl^ M(IL-4) (Fig. S4f, g), indicating that *Vavi*Cre x *Cpt1a*^fl/fl^ M(IL-4) are able to utilze other carbon sources to maintain glutamate for glutathione synthesis. Consistent with these data, we found significantly increased ROS levels in *Cpt1a*-shRNA and ERT2Cre x *Cpt1a*^fl/fl^ BMDM, but not in *Vavi*Cre x *Cpt1a*^fl/fl^ BMDM, compared to in respective wildtype controls (Fig. 4i, Fig. S4h, i).

Taken together, these data indicate that inhibition of Lc-FAO due to loss of function of Cpt1a in IL-4 stimulated ERT2Cre x *Cpt1a*^fl/fl^ BMDM and *Cpt1a-*shRNA BMDM results in diminished cellular availability of glutamate to fulfil multiple requirements, including fueling the TCA cycle, and glutathione synthesis. These effects were not apparent in IL-4 stimulated *Vavi*Cre x *Cpt1a*^fl/fl^ BMDM in which, in contrast to in ERT2Cre x *Cpt1a*^fl/fl^ BMDM and *Cpt1a-*shRNA BMDM, glutamine metabolism was upregulated to maintain TCA cycle intermediates and glutathione pools. Downstream effects of these changes in *Cpt1a-*shRNA BMDM include reduced chromatin accessibility, and the reduction of the NRF2 oxidative stress response accompanied by increased ROS levels.

### Acute loss of Cpt1a dysregulates glutathione metabolism

Our data show that glutamate levels, and the levels of metabolites that require glutamate for synthesis, are diminished in BMDM in which Cpt1a loss of function is associated with inhibition of alternative activation. In addition to being generated intracellularly through the TCA cycle or glutaminolysis, glutamate can be taken up from the extracellular space, although this is believed to occur mainly in neurons and astroglia^27^. We nevertheless asked whether extracellular glutamate supplementation could restore alternative activation. Instead, we found that extracellular glutamate inhibited expression of ARG1, CD301 and RELMα in IL-4 stimulated WT BMDM (Fig. 5a). The inhibitory effects on RELMα expression were more marked in WT cells than in *Cpt1a*-deficient cells. Extracellular glutamate is a known inhibitor of x_c_-, a cystine/glutamate antiporter comprised of Slc7a11 and Slc3a2, in which glutamate export is linked to cystine uptake^28,29^, and consistent with this we found that exogenous glutamate limited cystine uptake in M(IL-4) (Fig. 5b). Erastin, an inhibitor of x_c_-, potently inhibited IL-4 induced alternative activation in both WT and ERT2Cre x *Cpt1a*^fl/fl^ BMDM (Fig. S5a), pointing to a critical role for glutamate export coupled to cystine import in M(IL-4).

**Figure 5.**
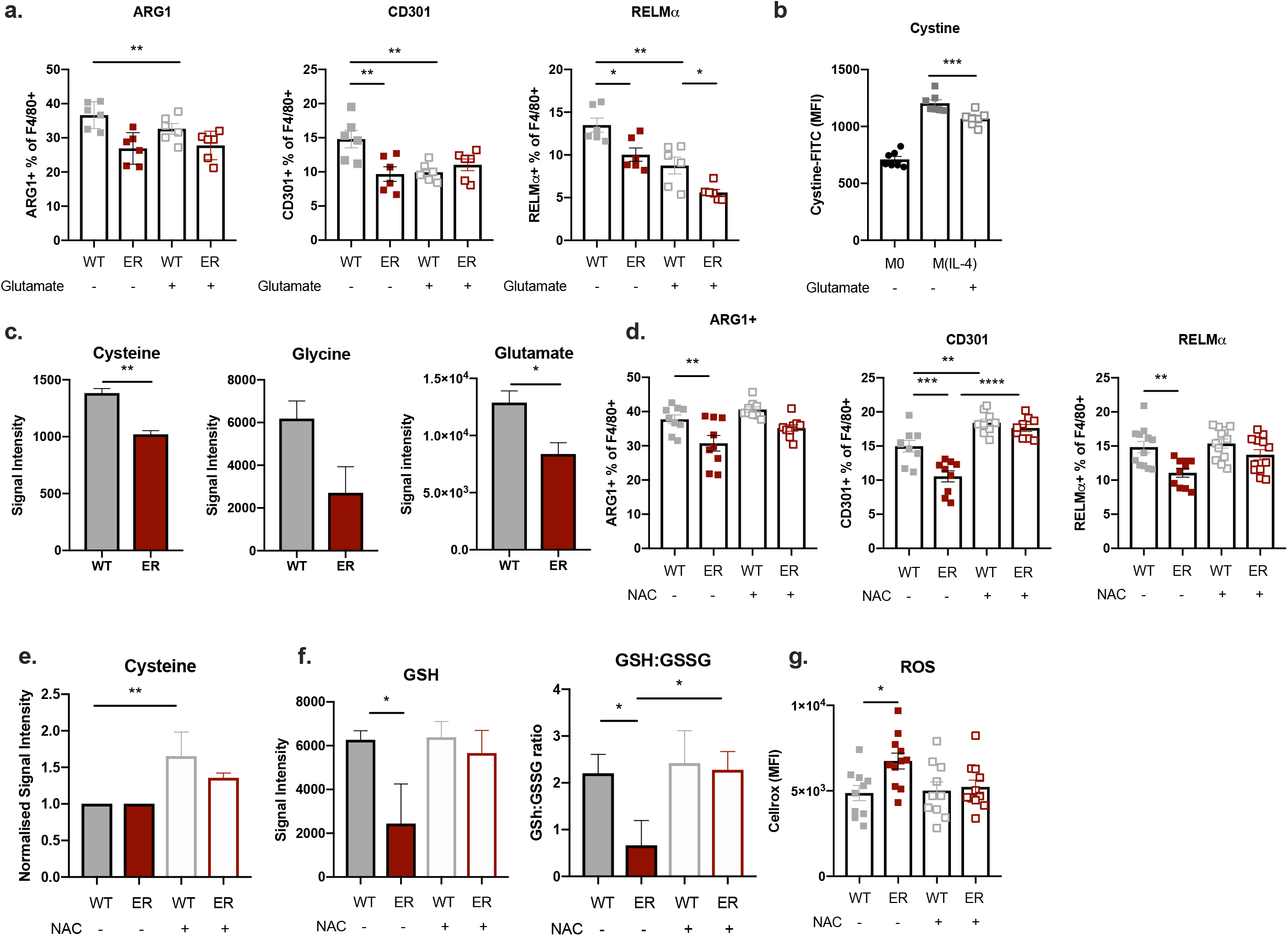
N-acetyl cysteine restores response to IL-4 in macrophage after acute loss of Cpt1a. a. Expression of ARG1, CD301 and RELMα as percentage of F4/80+ cells in *Cpt1a*^fl/fl^ (WT, grey) or ERT2Cre x *Cpt1a*^fl/fl^ (ER, red) IL-4 stimulated (M(IL-4)) BMDM cultured without (-) or with (+) added glutamate, as measured by flow cytometry. b. Median fluorescence intensity (MFI), as measured by flow cytometry, of intracellular FITC labeled cystine added for 5 min in cystine free media to resting M0 or M(IL-4) BMDC, without or with added glutamate, as indicated. c. Intracellular levels of cysteine, glycine and glutamate in WT and ER M(IL-4), as measured by LC-MS. d. Expression of ARG1, CD301 and RELMα as percentage of F4/80+ cells in WT or ER M(IL-4) BMDM cultured +/−NAC, as measured by flow cytometry. e. Intracellular levels of cysteine in WT and ER M(IL-4) treated +/−NAC, as measured by GC-MS and LC-MS. f. Intracellular levels of reduced glutathione (GSH) and the ratio of reduced to oxidized glutathione (GSH:GSSG) in WT and ER M(IL-4) treated +/− NAC, as measured by GC-MS and LC-MS. g. Cellular reactive oxygen species (ROS) determined by cellrox median fluoresence intensity (MFI) of WT and ER M(IL-4) BMDM treated +/− NAC, as measured by flow cytometry. Data represent mean ± SEM represent mean ± SEM from 6 (a, b) and 12 (d, e) biological replicates. Data represent three (a, c, e, f) and two (b, d) independent experiments. (*p<0.05, **p < 0.01, ***p<0.001, ****p < 0.0001).

Glutathione is synthesized from glutamate, cysteine and glycine. Our data suggested that cells in which glutamate is limiting might be challenged to take up sufficient cystine to meet glutathione needs. To examine this, we measured cysteine in IL-4 stimulated ERT2Cre x *Cpt1a*^fl/fl^ BMDM and WT BMDM, and found that loss of Cpt1a function was associated with reduced cellular pools of this amino acid, and parallel reductions in cellular pools of glycine and glutamate (Fig. 5c), similar to the effect in *Cpt1a*-shRNA compared to EV M(IL-4) (Fig. 4e). In contrast, cystine and glycine levels in IL-4 stimulated *Vavi*Cre x *Cpt1a*^fl/fl^ BMDM were similar to those in WT BMDM (Fig. S5b). In addition to acquiring exogenous cystine, BMDM can synthesize cysteine and glycine from glucose *via* serine. However, in line with the downregulation of genes involved in this pathway (Fig. 4g), ^13^C glucose contribution to serine and glycine was significantly diminished in control and IL-4-stimulated *Cpt1a*-shRNA and *Cpt2* BMDM compared to in EV-shRNA BMDM (Fig. S5c).

Finally, we reasoned that if decreased glutathione synthesis is linked to diminished alternative activation in macrophages deficient in Cpt1a, then restoring cysteine pools should be able to rescue the expression of key marks of alternative activation and reduced ROS to WT levels in parallel. Consistent with this ARG1, CD301 and RELMα expression in IL-4 stimulated ERT2Cre x *Cpt1a*^fl/fl^ BMDM were returned to WT levels by the addition of N-acetyl-cysteine (NAC) (Fig. 5d). NAC supplementation increased intracellular cysteine pools (Fig. 5e) and restored intracellular glutathione pools, which in turn increased the GSH:GSSG ratio in ERT2Cre x *Cpt1a*^fl/fl^ M(IL-4) (Fig. 5f), which reduced oxidative stress (Fig. 5g). Taken together our data suggest that Cpt1a expression is necessary to maintain glutamate levels to fuel glutathione synthesis and avoid ROS accumulation in M(IL-4). NAC supplementation during activation increases cysteine pools and restores glutathione pools, alleviating oxidative stress and allowing WT levels of expression of alternative activation markers.

## Discussion

We have found that, depending on the method used to target Cpt1a, loss of function of this enzyme can result in the inhibition of IL-4-induced alternative activation. Expression of M(IL-4) markers ARG1, CD301 and RELMα was impaired in ERT2Cre x *Cpt1a*^*fl/fl*^ macrophages and in macrophages expressing shRNA targeting *Cpt1a (Cpt1a-*shRNA). In contrast, consistent with previous reports^16,18^, expression of alternative activation marks in IL-4 stimulated *Lysm*Cre x *Cpt1a*^*fl/fl*^ macrophages was indistinguishable from in WT BMDM. We found that *Vavi*Cre x *Cpt1a*^*fl/fl*^ macrophages behaved similarly to *Lysm*Cre x *Cpt1a*^*fl/fl*^ macrophages, even though the degree of deletion varied. Our findings show that, in certain cases, macrophages are able to compensate for disruption of one metabolic pathway by re-routing other metabolic pathways to meet metabolic needs associated with the alternative activation program.

Our data show that reduced alternative activation due to Cpt1a loss of function was linked to reductions in cellular pools of glutamate. Consistent with previous reports^7,20^, we found that, glutamine metabolism is critical for alternative activation. Our data indicate that increased FAO in M(IL-4) serves to spare glutamate for use in other metabolic pathways that are critical for alternative activation. Amongst these pathways are αKG and glutathione production (Fig. 4a, c, d). αKG is a cofactor for Jmjd3-dependent demethylation of the repressive H3K27me3 mark associated with IL-4 responsive genes and as such has been previously reported to play a critical role in alternative activation^20^. Broadly consistent with this, ATAC-seq analysis revealed a marked reduction in accessibility around transcriptional start sites in *Cpt1a*-deficient M(IL-4). Nevertheless, we were unable to restore full alternative activation in ERT2Cre x *Cpt1a*^*fl/fl*^ or *Cpt1a-*shRNA M(IL-4) with supplemental αKG or cell-permeable dimethyl αKG (data not shown). Glutathione is important for the detoxification of ROS^25^. The addition of N-acetyl cysteine allowed the restoration of glutathione pools and of alternative activation in the face of *Cpt1a*-deficiency, pointing to a role for FAO in sparing glutamate for glutathione synthesis and the control of ROS.

ROS production is a by-product of mitochondrial respiration, and increased reliance on mitochondrial respiration, as in IL-4 stimulated macrophages, would be expected to lead to the need for enhanced ROS detoxification methods. Our previous work showed that alternatively activated macrophages have significantly lower cellular ROS levels than their inflammatory counterparts^30^. We hypothesize that this reflects sufficiency of glutamate for glutathione synthesis to alleviate oxidative stress. Glutathione is synthesized from glutamate, glycine and cysteine. We found that glycine biosynthesis from glucose carbon and cystine uptake were increased in IL-4 stimulated macrophages, but diminished when Cpt1a loss of function caused a reduction in alternative activation. Indeed, we found that increased ROS associated with reduced alternative activation due to Cpt1a loss of function, was associated with decreased expression of genes encoding proteins in the serine/glycine synthesis pathway, and more generally with a decrease in the expression of genes in the NRF2 stress response which is critical for the control of oxidtative stress (Fig. 4c, d).

We found that erastin, an inhibitor of x_c_-, strongly inhibits alternative activation. x_c_-is an amino acid transporter made up of Slc7a11 and Slc3a2, that couples the export of glutamate to the import of cystine^28,29^. The importation of cystine, which is required for glutathione synthesis, therefore requires adequate glutamate reserves to allow Slc7a11 activity. These findings support the view that alternative activation require the use of pathways that support the maintenance of adequate glutathione pools for ROS detoxification. However, our data do not exclude the possibility that Slc7a11 plays additional roles in alternative activation because cysteine is involved in the synthesis of CoA, which is an important co-factor facilitating FAO^31^, and the provision of sulfur for iron-cluster synthesis, which is critical for electron transport chain function and therefore respiration^32^.

Our data showed that ERT2Cre-mediated deletion of Cpt1a reduced the ability of IL-4-activated BMDM to promote B16 melanoma growth. This is consistent with prior work which showed that CD36-deficient macrophages, in which FAO would be expected to be inhibited, or etomoxir treatment *in vivo*, impair tumour growth^33^. The role of macrophages in immunity to cancer is complex. Inflammatory macrophages are implicated in the control of tumor growth and metastasis, but resting or alternatively activated macrophages can have the opposite effects and promote cancer progression^1,6^. However, the molecular cellular basis for tumor promotion by alternatively activated macrophages is unclear, although it may be related to immunosuppression resulting from arginine consumption by ARG1^34^. We found that macrophages in which Cpt1a was deleted by ERT2cre, expressed higher basal levels of markers of inflammatory activation, such as CD86 and MHC class II (data not shown), which could explain the trend, in our experiments, for Cpt1a-deficient macrophages, in comparison to WT macrophages, to limit tumor growth.

Our findings revealed a failure of *Cpt1a*-deficiency to affect alternative activation in *Vavi*Cre x *Cpt1a*^fl/fl^ and *Lysm*Cre x *Cpt1a*^fl/fl^ macrophages, which is consistent with earlier reports^18^. We found that an underlying difference between *Vavi*Cre x *Cpt1a*^fl/fl^ M(IL-4) versus ERT2Cre x *Cpt1a*^fl/fl^ and *Cpt1a*-shRNA, in which alternative activation is impaired, is that glutamine metabolism is upregulated in *Vavi*Cre x *Cpt1a*^fl/fl^ cells, and that this allows maintenance of αKG and glutathione pools and control of ROS, with minimal effects on bioenergetics. We reason that these changes are sufficient to support alternative activation. At present the underlying mechanisms that account for the differences in glutamine metabolism between *Vavi*Cre x *Cpt1a*^fl/fl^ M(IL-4) and ERT2Cre x *Cpt1a*^fl/fl^ or *Cpt1a*-shRNA M(IL-4) are unclear. Initially we hypothesized that this might reflect relative differences in deletion of *Cpt1a*, however Cpt1a is undetectable in *Vavi*Cre x *Cpt1a*^fl/+^ macrophages, yet these cells were able to up-regulate ARG1, CD301 and RELMα to the same extent as WT cells in response to IL-4. We postulate that differences in the effect of the loss fo Cpt1a in *Vavi*Cre x *Cpt1a*^fl/fl^ versus ERT2Cre x *Cpt1a*^fl/fl^ and *Cpt1a*-shRNA M(IL-4) may reflect differences in cellular adaptation to chronic Cpt1a deficiency (*Vavi*Cre x *Cpt1a*^fl/fl^, *Lysm*Cre x x *Cpt1a*^fl/fl^) versus more acute loss of function (ERT2Cre x *Cpt1a*^fl/fl^, *Cpt1a*-shRNA). Our findings that acute inhibition of MPC was able to inhibit alternative activation, whereas macrophages in which MPC was deleted by *Lysm*Cre-were able to become alternatively activated in response to IL-4, and that glutamine metabolism was enhanced in the latter cells to compensate for diminished glucose oxidation, supports the idea that macrophages are able to compensate for chronic disruption of one metabolic pathway by re-routing other metabolic pathways to meet metabolic needs. Our data raise questions about the blanket use of genetic loss of function models to draw conclusions about the importance of metabolic pathways for specific cellular functions.

Together, our data indicate that the coincident use of FAO and glutaminolysis to fuel OXPHOS can spare glutamate for the synthesis of glutathione, which is critical to regulate ROS in strongly respiring cells, like IL-4 stimulated macrophages. Nevertheless, under specifc circumstances, these cells can retain glutathione levels and the ability to become alternatively activated independently of FAO. This flexibility is linked to adaptive increases in glutamine metabolism.

## Acknowledgements

The authors thank Jörg Büscher and the Metabolomics Core at the MPI-IE for their advice and support of this project, and Andrea Quintana and John Sutherland for help with animal colonies. The work was supported by German Research Foundation (DFG) FOR 2599 (EJP), NIH AI 110481 (EJP), DFG under Germany’s Excellence Strategy (CIBSS EXC-2189 Project ID 390939984), the Alexander von Humboldt Fellowship Foundation (AK), the CAPES/Alexander von Humboldt Fellowship Foundation (88881.136065/2017-01) (AC), Japan Society for the Promotion of Science (MM), the Sao Paulo research foundation-FAPESP (2016/23328-0) (LBM), and the Max Planck Society.

## Materials & Methods

### Mouse models and *in vivo* experiments

C57BL/6J (RRID: IMSR_JAX:000664) mouse strain was purchased from The Jackson Laboratory. *Lysm*Cre x *Cpt1a*^*fl/fl*^, *Vavi*Cre x *Cpt1a*^*fl/fl*^ and ERT2Cre x *Cpt1a*^*fl/fl*^ mice were generated by crossing *Cpt1a*^fl/fl^ mice^35^ with *Lysm*Cre (JAX 004781), VaviCre (JAX 008610) or ERT2Cre (JAX no) mice respectively, while maintaining them on a C57BL/6 genetic background. *Lysm*Cre x *Mpc2*^*fl/fl*^ were generated by crossing *Mpc2*^fl/fl^ mice (Brian N. Finck, Washington University School of Medicine, St. Louis) to *Lysm*Cre mice. All mouse strains were maintained at the Max Planck Institute for Immunobiology and Epigenetics and cared for according to the Institutional Animal Use and Care Guidelines with approval by the animal care committee of the Regierungspraesidium Freiburg, Germany. All animals used for tissue harvest or experimental procedures were either male or female, aged between 6-12 weeks at the start of the experiment. Animals were humanely sacrificed by carbon dioxide asphyxiation followed by cervical dislocation and bone marrow or peritoneal lavage was harvested. Mice were bred under specific pathogen free standards.

ERT2Cre x *Cpt1a*^*fl/fl*^ mice were gavaged with 100µL 20mg/mL tamoxifen (Sigma) in sunfloweroil (Sigma) every day for 5 days after which the mice were sacrificed on the to collect the peritoneal lavage.

C57BL/6J mice were co-injected with 1×10^5^ WT-F10-B16-melanoma (ATCC) and 1×10^5^ pre-stimulated ERT2Cre x *Cpt1a*^*fl/fl*^ or wildtype M0 or M(IL-4). Tumour bearing mice were weighed and the tumour diameter measured every 2 days from day 3 after injection onwards. Mouse body condition was monitored over the experimental period to ensure that the mice maintained expected weight, did not show signs of distress, tumours did not ulcerate, impair movement or exceeded 20 mm.

### Primary cell cultures

Bone marrow cells were grown in complete medium (RPMI-1640 medium containing 10 mM glucose, 2 mM L-glutamine, 100 U ml^−1^ penicillin-streptomycin and 10% FCS) with 20 ng ml^−1^ murine macrophage colony-stimulating factor 1 (CSF-1; Peprotech) for 7 days, and supplemented with CSF-1 on days 2 and 4. Cre-ER expression was induced with 1µM 4-OHT (Sigma) from the start of the BMDM culture. On day 7 macrophages were harvested and then maintained in 20 ng ml^−1^ CSF-1 for subsequent experiments in which they were either maintained in medium without any further additions (M0) or with addition of 20 ng ml^−1^ IL-4 (Peprotech; MI(L-4)), for 20 h. M(IL-4)s could additionally be supplemented with 100µM palmitate, 5µM BPTES, 50µM UK5099, 1mM glutamate, 10µM Erastin or 1mM N-Acetyl Cysteine, (all Sigma).

Peritoneal macrophages were obtained by performing a peritoneal lavage on naive animals with 2% fetal bovine serum (GIBCO) in PBS. Cells were counted, plated in complete RPMI supplemented with CSF-1 and selected based on adherence. Stimulation was performed using the same conditions as with BMDM.

### Lentiviral production and cell transduction

HEK293T cells were transfected using Lipofectamine 3000 (Thermo Fisher Scientific) with lentiviral packaging vectors pCAG-eco and psPAX.2 plus empty pLKO.1 control (EV) with a puromycin selection cassette (all obtained from Addgene) or a shRNA containing pLKO.1 targeting Cpt1a (GE Dharmacon CAT# RMM3981-201819967 – TRCN0000110596) or Cpt2 (GE Dharmacon CAT# RMM3981-201824060 – TRCN0000110500). Virus was collected from the supernatant of the cells. Bone marrow cultures were transduced in the presence of polybrene (8 mg ml–1) on day 2 of culture. A 48 h, selection of transduced cells was performed with 6 μg ml^−1^ puromycin (Sigma).

### Flow cytometry

Used fluorochrome-conjugate monoclonal antibodies included: CD301 (Milteny Biotech, clone: REA687), CD11b (Biolegend, clone: M1/70), F4/80 (Biozol, clone: BM8), TIM4 (BioLegend, clone: F31-5G3). Staining was performed in 1% fetal bovine serum in PBS with 2mM EDTA for 20 min at 4 °C, including anti-CD16/CD32 (Biozol); dead cells were excluded with the LIVE/DEAD Fixable Blue Dead Cell Stain Kit (Thermo scientific). For intracellular ARG1 and RELMα staining, cells were processed with the Transcription Factor Staining Buffer Set (Thermo Fisher Scientific) following the manufacturer protocol, after which straining for ARG1 antibody staining (eBioscience, clone: A1exF5) and RELMα primary antibody staining (Peprotech Cat# 500-P214) proceeded. RELMα was detected using Alexa Fluor-421 anti-rabbit secondary antibody (Life technologies). For cellular ROS staining, macrophages were incubated with 50 μM Cellrox Deep Red (Thermo Fischer) in complete RPMI for 30 min at 37 °C. Cells were analyzed using LSR Fortessa flow cytometers (BDBiosciences) and data were processed using FlowJo software (FlowJo).

### Gene expression analysis by quantitative PCR with reverse transcription

RNA was isolated using RNAsolv (Omega Bio-Tek) and single-strand cDNA was synthesized using the High Capacity cDNA Reverse Transcription Kit (Applied Biosystems). Cpt1a (Mm01231182 or Mm01231183), Cpt1b (Mm01308154_m1), Cpt1c (Mm00463970) and Cpt2 (Mm00487205) RT-PCR was performed with Taqman primers using an Applied Biosystems 7000 sequence detection system. The expression levels of mRNA were normalized to the expression of HPRT.

### Western Blot

For western blot analysis, cells were washed with ice cold PBS and lysed at a concentration of 25,000 cell/μL and loaded protein from 175,000 cells/lane in 1 x Cell Signaling lysis buffer (20 mM Tris-HCl, [pH 7.5], 150 mM NaCl, 1 mM Na_2_EDTA, 1 mM EGTA, 1% Triton X-100, 2.5 mM sodium pyrophosphate, 1 mM β-glycerophosphate, 1 mM Na_3_VO_4_, 1 μg/mL leupeptin (Cell Signaling Technologies), supplemented with 1 mM PMSF. Samples were frozen and thawed 3 times followed by centrifugation at 20,000 x g for 10 min at 4°C. Cleared protein lysate was denatured with LDS loading buffer for 10 min at 70°C, and loaded on precast 4% to 12% bis-tris protein gels (Life Technologies). Proteins were transferred onto nitrocellulose membranes using the iBLOT 2 system (Life Technologies) following the manufacturer’s protocols. Membranes were blocked with 5% w/v milk and 0.1% Tween-20 in TBS and incubated with the appropriate antibodies in 5% w/v BSA in TBS with 0.1% Tween-20 overnight at 4°C. All primary antibody incubations were followed by incubation with secondary HRP-conjugated antibody (Pierce) in 5% milk and 0.1% Tween-20 in TBS and visualized using SuperSignal West Pico or femto Chemiluminescent Substrate (Pierce) on Biomax MR film (Kodak). Cpt1a antibody (ProteinTech, 15184-1-AP and Abcam, 8F6AE9), Cpt2 antibody (ProteinTech, 26555-1).

### ^13^C metabolite tracing

Cells (1×10^6^) were cultured in RPMI 1640 supplemented with 100µM ^13^C-palmitate, RPMI 1640-glucose supplemented with 10mM ^13^C-Glucose or RPMI 1640-Glutamine supplemented with 2mM ^13^C-Glutamine for 20 h, after which they were rinsed with cold 0.9% NaCl and extracted using 50:30:20 methanol:acetonitrile:H2O pre-cooled at −80 °C, and dried using a EZ-2 Elite evaporator (Genevac). Label tracing was carried out using an Agilent 1290 Infinity II UHPLC inline with a Bruker impact II QTOF-MS operated in full scan (MS1) mode or gas chromatography mass spectrometry (Agilent 5977). Data processing including correction for natural isotope abundance was performed by an in-house R script. Metabolite peaks were identified based on exact mass and matching of retention time to a pure standard, correction for natural isotope abundance and calculation of fractional contribution was performed as described elsewhere.^36^

### Seahorse Extracellular Flux Assays

Extracellular acidification rate (ECAR) and oxygen consumption rate (OCR) were measured using the 96 well Extracellular Flux Analyzer (Seahorse Bioscience). 1×10^5^ BMDM exposed to different treatments were plated each well of seahorse XF96 cell culture and preincubated at 37°C for a minimum of 45 min in the presence of the same treatments, in the absence of CO_2_ in un-buffered RPMI with 10 mM glucose, with or without 2 mM L-glutamine, with pH adjusted to 7.4. OCR and ECAR were measured under basal conditions, after activation, and after the addition of the following drugs: 1 μM oligomycin, 1.5 μM flurorcarbonyl cyanide phenylhydrazon (FCCP) and 100 nM rotenone + 1 μM antimycin A (all SIGMA) as indicated. Results were collected with Wave software version 2.4 (Agilent).

### Metabolite quantification with Bioanalyzer

100 μL overnight culture media of 100.000 cells/200 μL was analyzed on the CEDEX Bio Analyzer (Roche), using the Glutamine V2 Bio or Glucose Bio test kits.

### ATAC-seq

Libraries were prepared using the Nextera DNA library Prep Kit (Illumina) adapting a published protocol.^37^ Briefly, 5×10^4^ BMDM treated for 20 hr as described were washed in PBS and then lysed in 10 mM Tris-HCl, pH 7.4,10 mM NaCl, 3 mM MgCl_2_ and 0.1% Igepal CA-630 (all SIGMA). Nuclei were then spun down and then resuspend in 25 μL TD (2x reaction buffer), 2.5 μL TDE1 (Nextera Tn5 Transposase) and 22.5 μL nuclease-free water, incubated for 30 min at 37°C. DNA was purified with the QIAGEN MinElute PCR Purification Kit (Thermo Fisher Scientific). PCR amplification was performed with the NEBNext High-Fidelity 2x PCR Master Mix (New England Labs) using custom Nextera PCR Primers containing barcodes. Adaptors were removed with AMPure XP beads according to manufacturer’s protocol. Libraries were quantified with the Qubit and submitted for sequencing with a HISeq 3000 (Illumina) by the staff at the Deep-sequencing Facility at the Max-Planck-Institute for Immunobiology and Epigenetics.

Sequenced samples were trimmed with Trimmomatic,^38^ mapped using Bowtie2^39^ and replicate mapped files merged with SAM tools.^40^ Coverage files were generated with deepTools.^41^ Open chromatin and differentially regulated chromatin was detected with MACS2^42^ with a p value < 1 × 10^−7^ and a q value of less than 0.1 and a 2-fold enrichment threshold. Bed files were analyzed with Bedtools,^43^ and visualized alongside coverage files on IGV.^44^

### RNA sequencing

Total RNA was extracted with RNAsolv (Omega Bio-Tek) and quantified using Qubit 2.0 (Thermo Fisher Scientific) according to the manufacturer’s instructions. Libraries were prepared using the TruSeq stranded mRNA kit (Illumina) and sequenced in a HISeq 3000 (Illumina) by the Deep Sequencing Facility at the Max Planck Institute for Immunobiology and Epigenetics. Sequenced libraries were processed with a pipeline optimized by the Bioinformatics core at the Max Planck Institute for Immunobiology and Epigenetics.^45^ Raw mapped reads were processed in R (Lucent Technologies) with DESeq2^46^ to determine differentially expressed genes and generate normalized read counts to visualize as heatmaps using Morpheus (Broad Institute).

### Statistical Analysis

Statistical analysis was performed using prism 6 software (Graph pad) and results are represented as mean ± SEM. Comparisons for two groups were calculated using unpaired two-tailed Student’s t tests, comparisons of more than two groups were calculated using one-way ANOVA with Sidak’s multiple comparison tests. We observed normal distribution and no difference in variance between groups in individual comparisons. Statistical significance: *p<0.05; ** p < 0.01; *** p < 0.001; **** p < 0.0001. Further details on statistical analysis are listed in the figure legends.

## Supplementary Figure Legends

**Supplementary figure 1.**
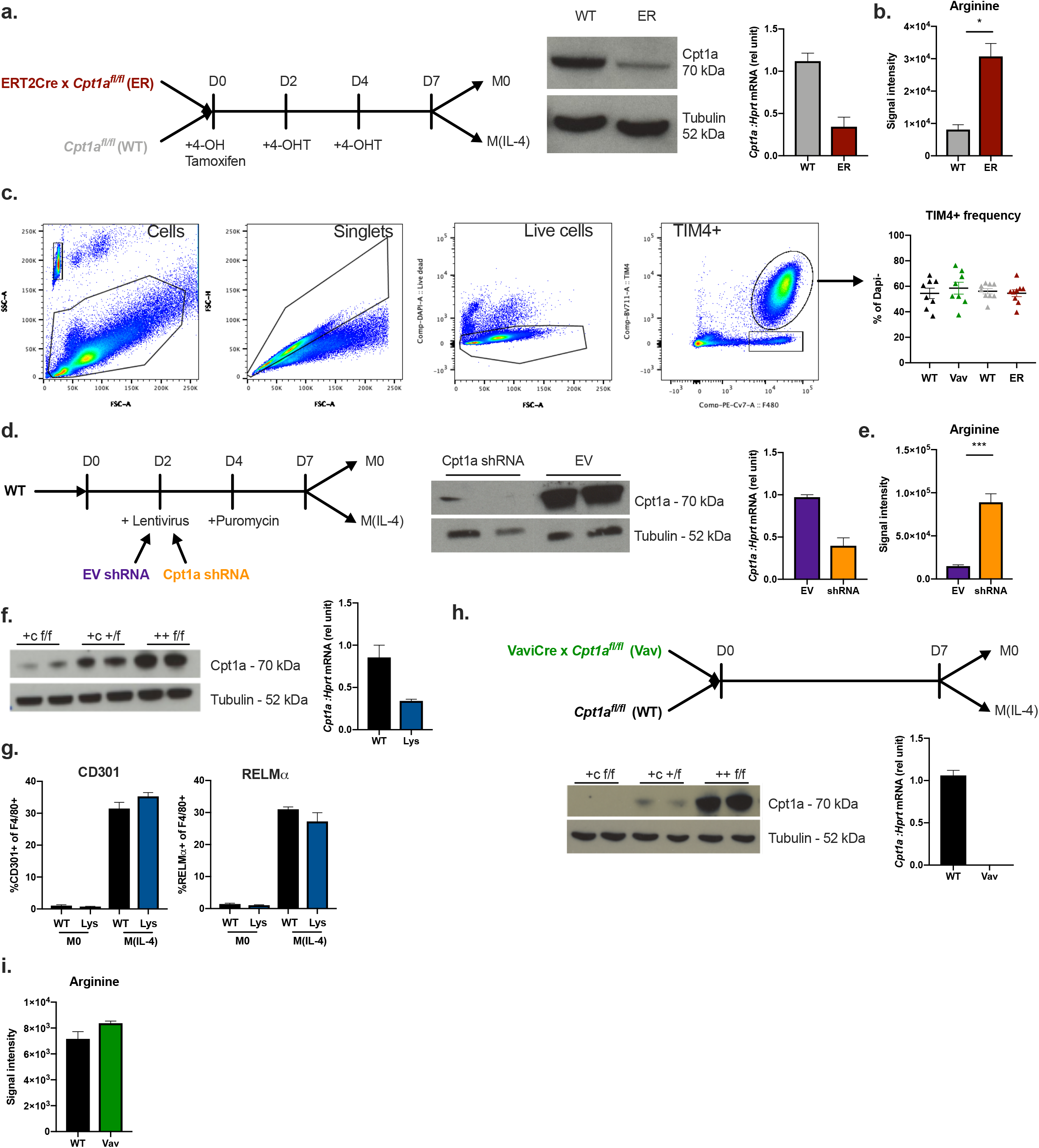
a ERT2Cre x *Cpt1a*^*fl/fl*^ (ER, red) or *Cpt1a*^*fl/fl*^ (WT, grey) BMDM were put in culture with 4-OH-Tamoxifen (1µM) added from day 0, and re-plenished during feeding on day 2 and day 4. On day 7 the cells were lifted from the plate and replated in the indicated conditions in resting (M0) or IL-4 stimulated (M(IL-4)) BMDM. Western blot of Cpt1a protein and q-PCR of *Cpt1a* mRNA on day 7 of culture. b. Intracellular arginine levels in ER or WT M(IL-4), as measured by LC-MS. c. Gating strategy of whole peritoneal lavage gating cells to singlets, to live cells (dapi-) to TIM4+F4/80+. TIM4+F4/80+ frequency in *Cpt1a*^*fl/fl*^ (WT, black) or *Vavi*Cre x *Cpt1a*^*fl/fl*^ (Vav, green) and WT (grey) or ER (red) of dapi-, live cells in whole peritoneal lavage samples. d. Wildtype BMDM were cultured and transduced with empty vector (EV, purple) or *Cpt1a*-shRNA (shRNA, orange) lentivirus on day 2 of culture. Cells were selected from day 4 of culture with puromycin and re-plated for stimulation on day 7. Western blot of Cpt1a protein and qPCR of *Cpt1a* mRNA on day 7 of culture. e. Intracellular arginine levels in EV or shRNA M(IL-4), as measured by LC-MS. f. Cpt1a protein expression as measured by western blot and mRNA expression as measured by qPCR in *Lysm*Crex *Cpt1a*^*fl/fl*^(+c f/f /*Lysm*Cre, blue), *Lysm*Cre x *Cpt1a*^*fl/+*^ (+c +/f) and *Cpt1a*^*fl/fl*^ (++f/f /WT, black) BMDMs. g. Expression of CD301 and RELMα as percentage of F4/80+ cells in WT and Lys resting (M0) or IL-4 stimulated BMDMs (M(IL-4)), as measured by flow cytometry. h. Vav BMDMs were put in culture and fed on day 2 and 4 of culture until stimulation on day 7. Cpt1a protein expression as measured by western blot in *Vavi*Cre x *Cpt1a*^*fl/fl*^ (+c/f/f), *Vavi*Cre x *Cpt1a*^*fl/+*^ (+c/+/f) and *Cpt1a*^*fl/fl*^ (++ f/f) BMDMs. Cpt1a mRNA expression as measured by qPCR in WT and Vav BMDMs, on day 7 of culture. i. Intracellular arginine levels in WT or Vav M(IL-4), as measured by LC-MS. Data represent mean ± SEM mean ± SEM from 3 (a, b), 9 (c), 4 (d, e, f, g, i), 2 (h biological replicates. Data represent three (a, d, f, g, h) and two (b, c, e, i) independent experiments. (*p<0.05, ***p<0.001).

**Supplementary figure 2.**
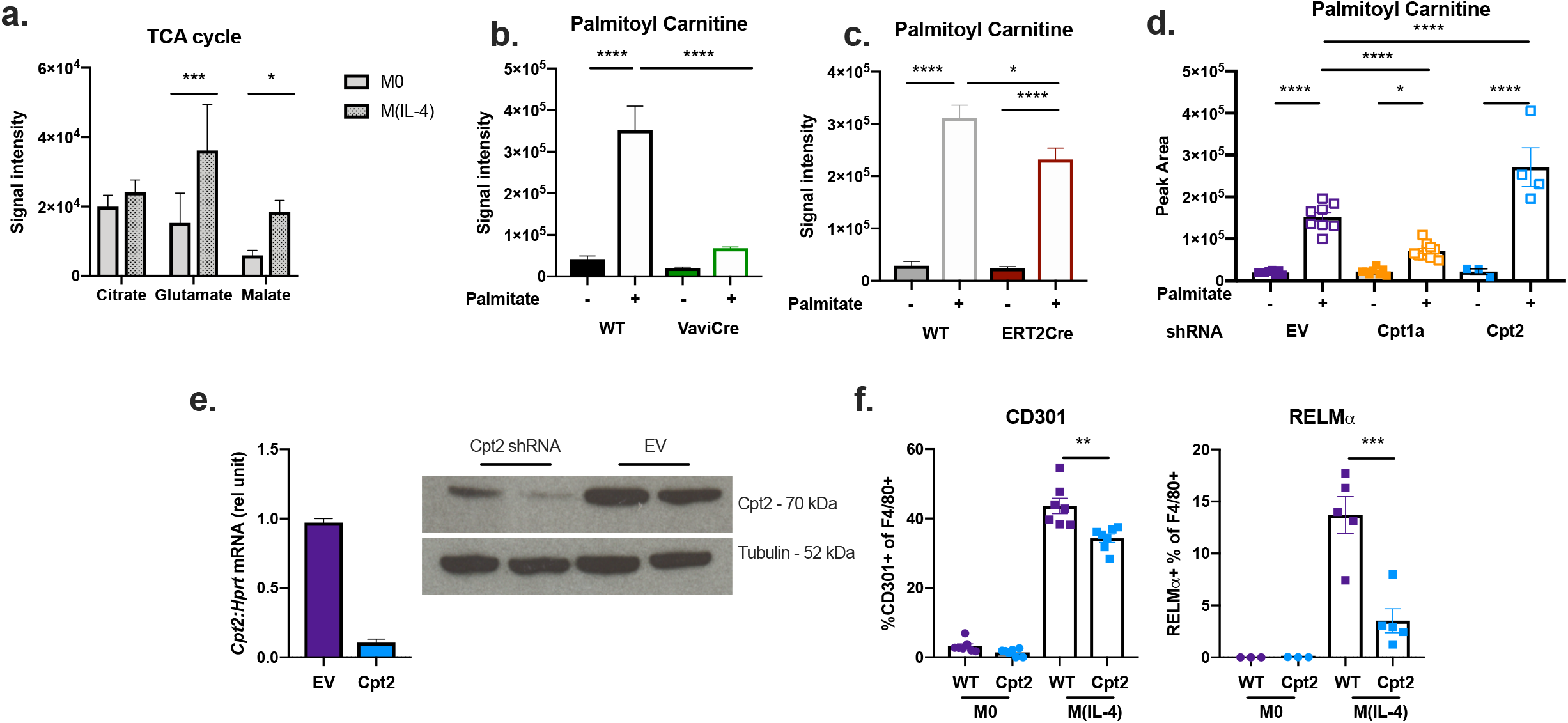
A Intracellular metabolite pools of citrate, glutamate and malate in wildtype resting (M0) or IL-4 stimulated BMDMs (M(IL-4)) as measured by LC-MS. b. Intracellular palmitoyl carnitine in *Cpt1a*^*fl/fl*^ (WT, black) or *Vavi*Cre x *Cpt1a*^*fl/fl*^ (Vav, green) M(IL-4) supplemented with (+) or without (-) 100 µM palmitate, as measured by LC-MS. c. Intracellular palmitoyl carnitine in *Cpt1a*^*fl/fl*^ (WT, grey) or ERT2Cre x *Cpt1a*^*fl/fl*^ (ER, red) supplemented +/−100 µM palmitate, as measured by LC-MS. d. Intracellular palmitoyl carnitine in empty vector (EV, purple), *Cpt1a-*shRNA (Cpt1a, orange) or *Cpt2-*shRNA (Cpt2, blue) M(IL-4) supplemented +/− 100 µM palmitate, as measured by LC-MS. *e. Cpt2* mRNA as measured by qPCR and Cpt2 protein as measured by western blot, in EV or *Cpt2*-shRNA BMDMs on day 7 of culture. f. Expression of CD301 and RELMα as percentage of F4/80+ cells in EV or Cpt2 M0 or M(IL-4), as measured by flow cytometry. Data represent mean ± SEM mean ± SEM from 7 (a), 4 (b), 3(c), 8/4 (d), 6 (e), 9 (f), 4 (g), 5 (h), biological replicates. Data represent three (a, e, f,h) and two (b, c, d, g) independent experiments. (*/#p<0.05, **/##p < 0.01, ***p<0.001, ****p < 0.0001).

**Supplementary figure 3.**
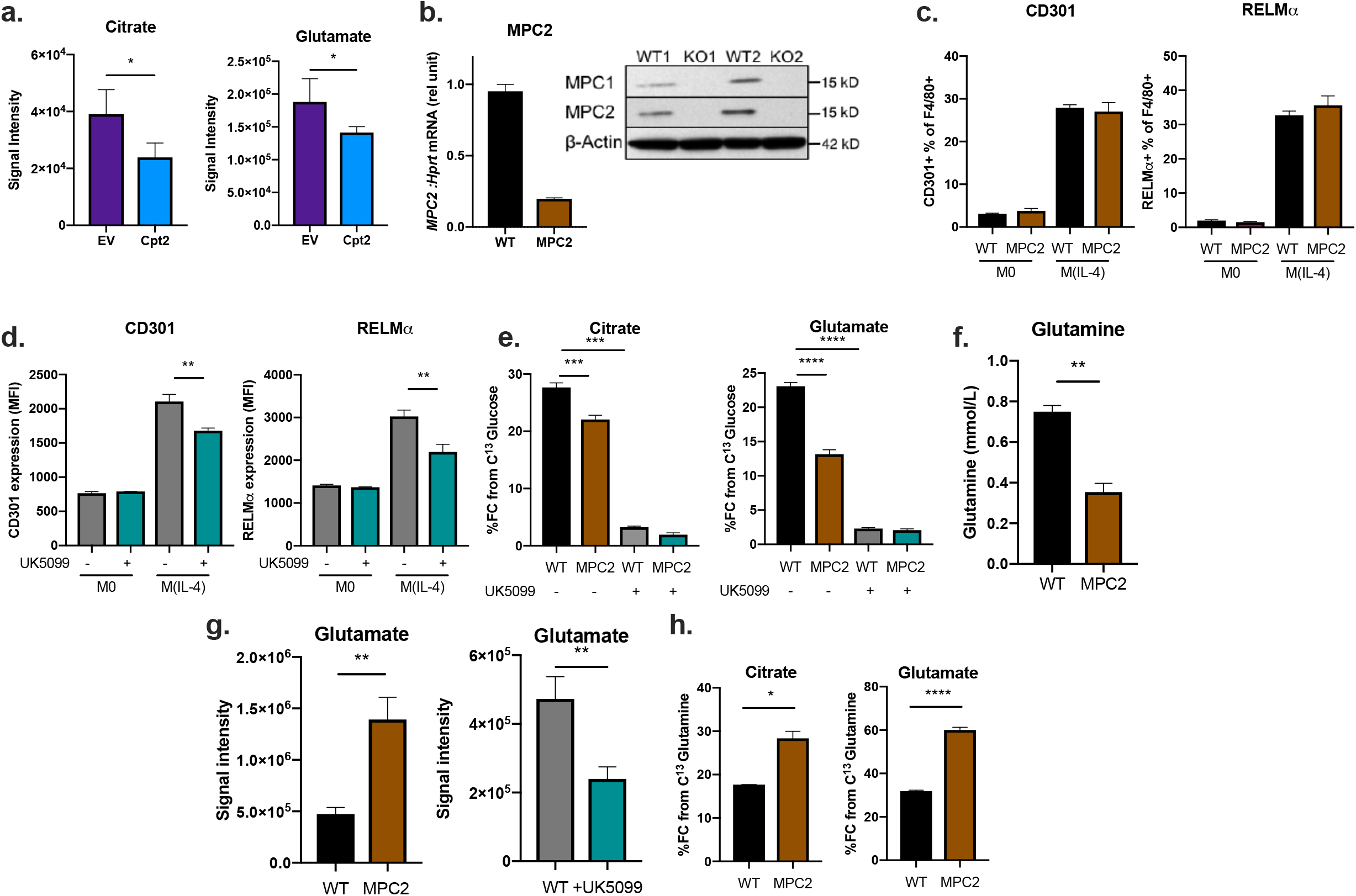
A,^13^C-Palmitate fractional contribution (FC) in citrate and glutamate in empty vector (EV, purple) or *Cpt2*-shRNA (Cpt2, blue) M(IL-4) as measured by LC-MS. b. *MPC2* mRNA as measured by qPCR and MPC1 and MPC2 protein expression as measured by western blot in *MPC2*^*fl/fl*^ (WT, black) or *Lysm*Cre x *MPC2*^*fl/fl*^ (MPC2, brown) BMDMs. c. Expression of CD301 and RELMα as percentage of F4/80+ cells in WT or MPC2 resting (M0) or IL-4 stimulated BMDMs (M(IL-4)), as measured by flow cytometry. d. Expression of CD301 and RELMα as percentage of F4/80+ cells in WT with (+, turqois) or without (-, grey) UK5099 M0 or M(IL-4), as measured by flow cytometry. e.. %^13^C-glucose fractional contribution (FC) contribution to TCA cycle intermediates citrate, glutamate and malate of WT or MPC2 M(IL-4) +/−UK5099, as measured by GC-MS. f. Glutamine concentration in mmol/liter in the supernatant of WT or MPC2 M(IL-4) as measured by bioanalyzer. g. Intracellular glutamate levels in WT or MPC2 and WT +/− UK5099, as measured by LC-MS. h. Fractional contribution (FC) from ^13^C Glutamine in citrate, glutamate and malate in WT or MPC2 M(IL-4) as measured by GC-MS. Data represent mean ± SEM mean ± SEM from 2 (a), 3 (b, c, d, e, f) or 4 (g) biological replicates. Data represent three (a, b, c, g) and two (d, e, f) independent experiments. (*p<0.05, **p < 0.01, ***p<0.001, ****p < 0.0001).

**Supplementary figure 4.**
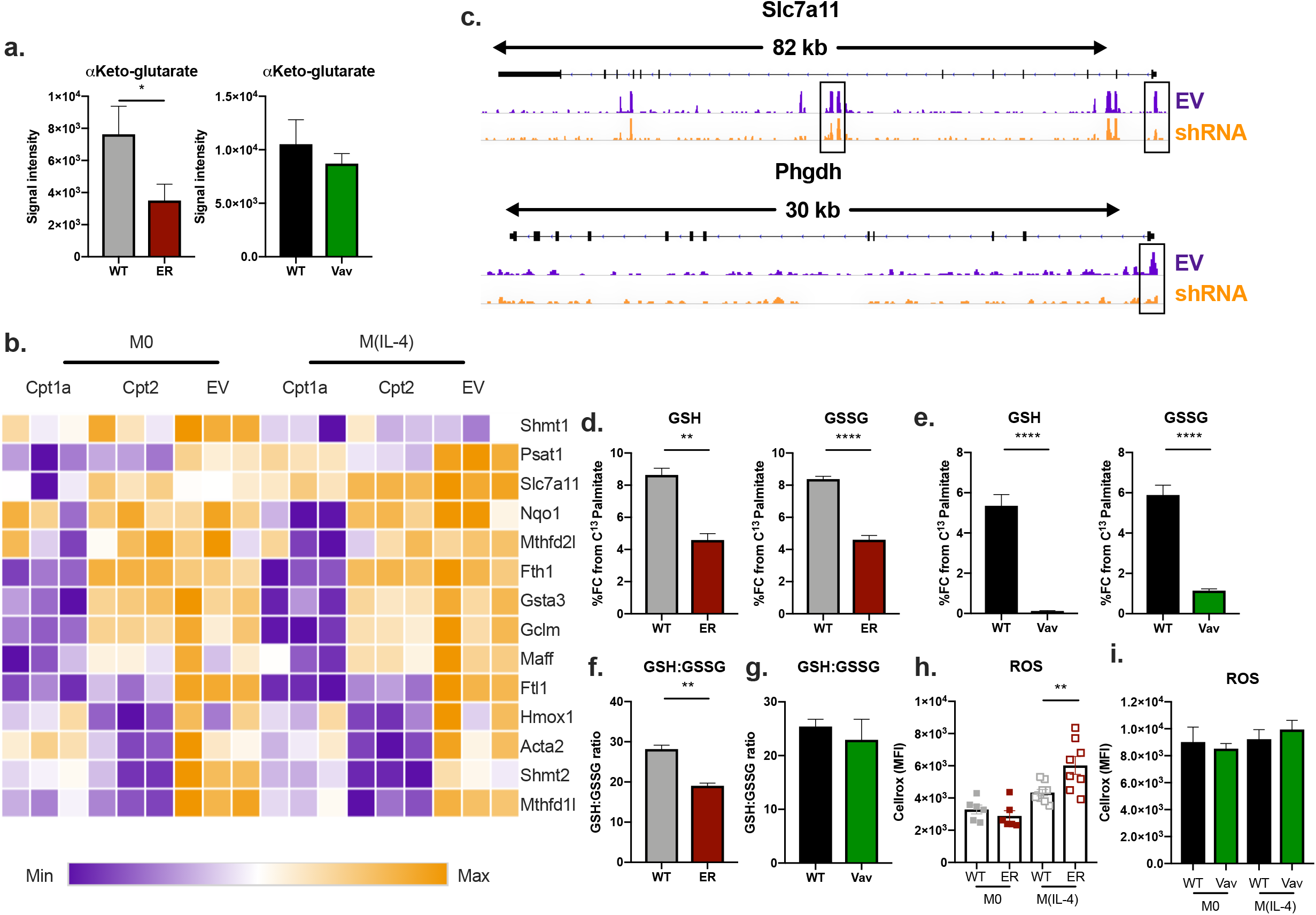
A Intracellular α-ketoglutarate levels, as measured by LC-MS, in *Cpt1a*^*fl/fl*^ (WT, grey) or ERT2Cre x *Cpt1a*^*fl/fl*^ (ER, red) and *Cpt1a*^*fl/fl*^ (WT, black) or *Vavi*Cre x *Cpt1a*^*fl/fl*^ (Vav, green) BMDMs stimulated with IL-4 (M(IL-4)). b. Heatmap of genes indicated as NRF2 oxidative stress response genes that were up/down-regulated significantly (p<0.01) more than 2-fold in Cpt1a shRNA M(IL-4) compared to EV M(IL-4). Heatmap visualizes log2fold change in M0 or M(IL-4) empty vector (EV), *Cpt1a-*shRNA (Cpt1a) or *Cpt2-*shRNA (Cpt2). c. Regions of accessible chromatin, measured by ATACseq, of *Slc7a11* and *Phgdh* genes in EV (purple track) and *Cpt1a*-shRNA (orange track) BMDMs stimulated with IL-4. The black boxes indicate regions of chromatin that are significantly less accessible in shRNA compared to EV. d. ^13^C-Palmitate contribution to reduced glutathione (GSH) and oxidized glutathione (GSSG) in WT or ER M(IL-4) as determined by LC-MS. e. ^13^C-Palmitate contribution to reduced glutathione (GSH) and oxidized glutathione (GSSG) in WT or Vav M(IL-4) as determined by LC-MS. f. Ratio fo GSH to GSSG signal intensities in WT or ER M(IL-4) as measured by LC-MS. g. Ratio fo GSH to GSSG signal intensities in WT or Vav M(IL-4) as measured by LC-MS. h. Median fluorescent intensity (MFI) of cellular ROS (cellrox) in WT or ER resting BMDMs (M0) or (M(IL-4)), as measured by flow cytometry. i. Median fluorescent intensity (MFI) of cellular ROS (cellrox) in WT or Vav M0 or (M(IL-4)), as measured by flow cytometry. Data represent mean ± SEM SEM ± standard deviation from 3 and 4 (a, b, c, d, e, f, h) or 10 (g) biological replicates. Data represent three (g, h) and two (a, b, c, d, e) independent experiments. (*p<0.05, **p < 0.01, ****p < 0.0001).

**Supplementary figure 5.**
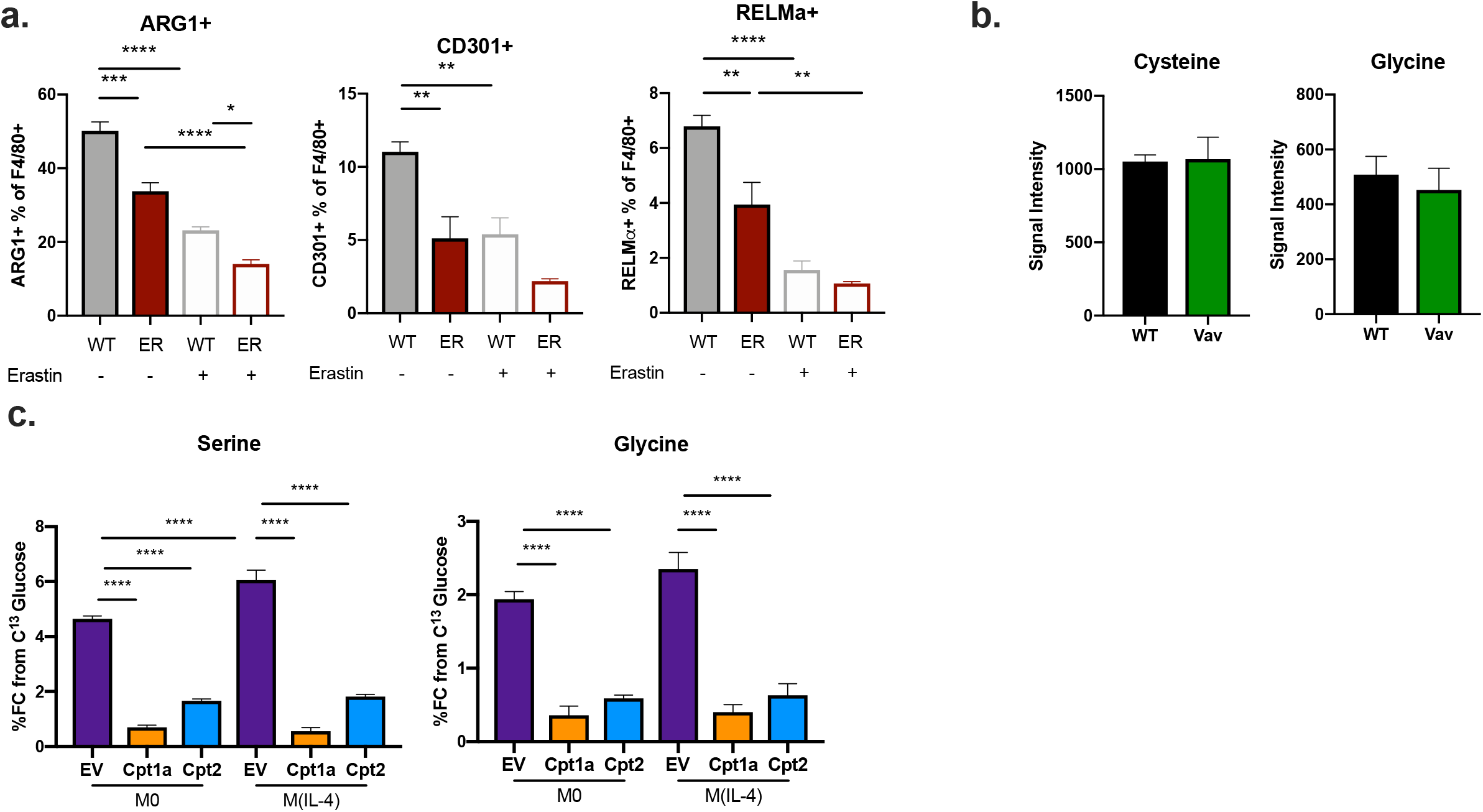
A Expression of Arginase1 (ARG1), CD301 and RELMα as percentage of F4/80+ cells in *Cpt1a*^fl/fl^ (WT, grey) or ERT2Cre x *Cpt1a*^fl/fl^ (ER, red) IL-4 stimulated (M(IL-4)) BMDM cultured without (-) or with (+) 10 µM erastin, as measured by flow cytometry. b. Intracellular cysteine and glycine levels in WT or Vav M(IL-4) as measured by LC-MS. c. ^13^C-glucose contribution to reduced serine and glycine in empty vector (EV), *Cpt1a-*shRNA (Cpt1a) or *Cpt2-*shRNA (Cpt2) in resting (M0) or IL-4 stimulated (M(IL-4)) BMDMs, as determined by GC-MS. Data represent mean ± SEM mean ± SEM from 4 biological replicates. Data represent three (a) and two (b, c) independent experiments. (*p<0.05, **p < 0.01, ****p < 0.0001).

